# Measuring the response to visually presented faces in the human lateral prefrontal cortex

**DOI:** 10.1101/2022.03.06.483119

**Authors:** Lara Nikel, Magdalena W Sliwinska, Emel Kucuk, Leslie G. Ungerleider, David Pitcher

## Abstract

Neuroimaging studies have identified multiple face-selective areas. In the current study we compared the functional response of the face area in the lateral prefrontal cortex to that of other face-selective areas. In Experiment 1 participants (N=32) were scanned viewing videos containing faces, bodies, scenes, objects, and scrambled objects. We identified a face-selective area in the right inferior frontal gyrus (rIFG). In Experiment 2 participants (N=24) viewed the videos or static images. Results showed that the rIFG, posterior superior temporal sulcus (pSTS) and occipital face area (OFA) exhibited a greater response to moving than static faces. In Experiment 3 participants (N=18) viewed face videos presented in the contralateral and ipsilateral visual fields. Results showed that the face areas in the IFG and pSTS responded equally to faces in both visual fields, while the OFA and fusiform face area (FFA) showed a contralateral bias. These experiments suggest two conclusions; firstly, in all three experiments the face area in the IFG was not as reliably identified as face areas in the occipitotemporal cortex. Secondly, the similarity of the response patterns in the IFG and pSTS face areas suggests that the areas are functionally connected, a conclusion consistent with neuroanatomical and functional connectivity evidence.

## Introduction

Faces are rich sources of social information that convey someone’s identity, attentional focus, and emotional state. Humans process this wealth of socially relevant information in a network of face-selective areas distributed across the brain (Calder & Young, 2005; Haxby et al., 2000). Three of the most heavily studied face-selective areas are in the occipitotemporal cortex and are thought to perform different cognitive functions. The fusiform face area (FFA) preferentially processes facial identity (Grill-Spector et al., 2004; Parvizi et al., 2012; Rezlescu et al., 2012; Rotshtein et al., 2005), the posterior superior temporal sulcus (pSTS) preferentially processes facial expressions (Winston et al., 2004; Pitcher, 2014) and the occipital face area (OFA) processes the component parts of the face (e.g., eyes and mouth) (Gauthier et al., 2000; Pitcher et al., 2007; Rossion et al., 2003). Beyond these core face-selective areas in visual cortex there is an extended network of additional face processing areas (Calder & Young, 2005; Haxby et al., 2000). One area identified in neural models of face processing is in the lateral prefrontal cortex. Studies of both humans and non-human primates report face-selective neural activity in the lateral prefrontal cortex (Haxby et al., 1996; Haxby et al., 1995; Ishai et al., 2002; Scalaidhe et al., 1997; Shepherd & Freiwald, 2018; Tsao et al., 2008) but how the lateral prefrontal cortex interacts with face-selective areas in the occipitotemporal cortex remains unclear. In the current study we compared the neural response to faces in the lateral prefrontal cortex with that observed in the more commonly studied face-selective areas in the occipitotemporal cortex.

Our prior knowledge and experience of the world shapes how we perceive incoming sensory input. The lateral prefrontal cortex is implicated in several neural processes that support these processes including cognitive control (MacDonald et al., 2000), working memory (Curtis & D’Esposito, 2003), and Theory of Mind (Kalbe et al., 2010). This range of different cognitive functions is consistent with evidence demonstrating that prefrontal areas are identified in face processing studies regardless of stimulus format, emotional valence, or task demands (Ishai et al., 2005). Neuroimaging studies of face processing have also demonstrated that the lateral prefrontal cortex is involved in the top-down control of ventral temporal cortex when recognising faces (Baldauf & Desimone, 2014; Heekeren et al., 2004). In addition, the lateral prefrontal cortex has been implicated in familiar face recognition (Rapcsak et al., 1996), working memory for faces (Courtney et al., 1996, 1997), famous-face recognition (Ishai et al., 2002), processing of information from the eyes (Chan & Downing, 2011) and configural processing of the component parts of faces (e.g. the eyes and mouth) (Renzi et al., 2013). Such a broad range of different face processing functions suggests that the connectivity between the lateral prefrontal cortex and other face processing areas may vary depending on the specific requirements of the current face processing task being performed.

The recognition of facial expressions of emotion is one of the functions processed in the lateral prefrontal cortex. Connectivity between the lateral prefrontal cortex and the amygdala has been demonstrated in healthy human participants (Davies-Thompson & Andrews, 2012) and this same circuit is thought to be impaired in mental illnesses such as major depressive disorder (MDD) (Heller et al., 2009). More recently a large-scale analysis of data collected from 680 participants reported a connection between the lateral prefrontal cortex and pSTS specialised for processing the dynamic facial aspects (Wang et al., 2020). The authors segregated the established nodes of the face processing network into three sub-networks using structural and functional connectivity analyses. Results demonstrated that the connectivity between the lateral prefrontal cortex and the pSTS was greater than between other face processing nodes. This is consistent with studies demonstrating that the pSTS preferentially processes dynamic facial aspects (Fox et al., 2009; Pitcher et al., 2019; Puce et al., 1998) and facial expressions (LaBar et al., 2003; Phillips et al., 1998; Sliwinska, Elson, et al., 2020; Winston et al., 2004). In addition, a study that assessed damage to the arcuate fasciculus (a white matter tract that connects the lateral temporal lobe with the inferior frontal lobe) reported behavioural impairments in face based mentalising tasks (Nakajima et al., 2018). These studies suggests that the lateral prefrontal cortex and pSTS may be nodes in a network for processing facial expressions, and particularly for processing the changes in faces that convey the emotions and intentions of other people.

The face-selective regions in the prefrontal cortex are also involved in accessing personal semantic information associated with a face. It has been suggested that they form part of a top-down sub-network which accesses existing knowledge associated with faces and is involved in decision making and working memory (Li et al., 2009). This is consistent with evidence showing that the inferior frontal gyrus (IFG) preferentially responds to famous faces, which, as opposed to recently learned faces, are processed beyond the stage of simple recognition to semantic identification (Ishai et al., 2005; Leveroni et al., 2000). The frontal activation may reflect long-term retrieval from a person-identity system by triggering and structuring the search for stored representations. Alternatively, the frontal regions may be part of view-independent face processing of familiar faces, as opposed to view-dependent processing of newly learned faces (Leveroni et al., 2000). This would mean that the frontal face area is involved in familiar face recognition without retrieving person-specific semantics. However, many studies support the involvement of the prefrontal areas not only in face processing but also in semantic retrieval.

Our aim was to better understand the functional connections between the face area in the lateral prefrontal cortex and the face-selective areas in the occipitotemporal cortex (namely, the OFA, FFA and pSTS). We did this by measuring the neural responses to different types of visual stimuli across the nominated face-selective regions of interest (ROIs) using functional magnetic resonance imaging (fMRI). In Experiment 1 we first established how robustly we could localise a face-selective neural response (defined using a contrast of faces greater than objects) in the lateral prefrontal cortex. We then compared the response to different categories of stimuli (faces, bodies, scenes, objects, and scrambled objects) in this area to that measured in the other face-selective areas. In Experiment 2 we measured the response to moving and static stimuli from these same visual categories across the face-selective areas. Prior studies have demonstrated that the pSTS exhibits a greater response to moving faces than static faces (Fox et al., 2009; Pitcher et al., 2019) but this same dissociation is not consistently observed in the FFA and OFA (Pilz et al., 2009; Pitcher et al., 2014; Schultz et al., 2013). Finally, in Experiment 3 we presented face videos depicting different facial expressions in the contralateral and ipsilateral visual fields. This was done to compare the visual field responses across the occipitotemporal face-selective areas with that of the face-selective area in the lateral prefrontal cortex. Prior studies have demonstrated that the contralateral visual field advantage observed in the FFA and OFA (Hemond et al., 2007; Kay et al., 2015) is absent in the STS (Finzi et al., 2021; Pitcher et al., 2020; Sliwinska, Bearpark, et al., 2020). We hypothesised that if the face areas in the pSTS and lateral prefrontal cortex are functionally connected then the lateral prefrontal cortex would also show an equal response to faces in both visual fields (thus distinguishing it from the FFA and OFA).

## Materials and Methods

### Participants

In Experiment 1 a total of thirty-two right-handed participants (18 females, 14 males) with normal, or corrected-to-normal, vision gave informed consent as directed by the Ethics committee at the University of York. In Experiment 2 twenty-four right-handed participants (17 females, 7 males) with normal, or corrected-to-normal, vision gave informed consent as directed by the Ethics committee at the University of York. In Experiment 3 eighteen participants (10 females, 8 males) with normal, or corrected-to-normal, vision gave informed consent as directed by the National Institutes of Mental Health (NIMH) Institutional Review Board (IRB). The data reported in Experiment 3 was collected for a previous visual field mapping experiment (Pitcher et al., 2020) and re-analysed for the current study.

### Stimuli

In all three experiments we used 3-second movie clips of faces and objects to localize the face-selective brain areas of interest (Pitcher et al., 2011; Pitcher et al., 2014). In Experiment 1 and 2 participants also viewed 3-second movie clips of bodies, scenes and scrambled objects to calculate the response profiles to different stimulus categories. There were sixty movie clips for each category in which distinct exemplars appeared multiple times. Movies of faces and bodies were filmed on a black background, and framed close-up to reveal only the faces or bodies of 7 children as they danced or played with toys or adults (who were out of frame). Fifteen different locations were used for the scene stimuli which were mostly pastoral scenes shot from a car window while driving slowly through leafy suburbs, along with some other films taken while flying through canyons or walking through tunnels that were included for variety. Fifteen different moving objects were selected that minimized any suggestion of animacy of the object itself or of a hidden actor pushing the object (these included mobiles, windup toys, toy planes and tractors, balls rolling down sloped inclines). Scrambled objects were constructed by dividing each object movie clip into a 15 by 15 box grid and spatially rearranging the location of each of the resulting movie frames. Within each block, stimuli were randomly selected from within the entire set for that stimulus category (faces, bodies, scenes, objects, scrambled objects). This meant that the same actor, scene or object could appear within the same block but given the number of stimuli this did not occur regularly.

In Experiment 2 static stimuli were identical in design to the dynamic stimuli except that in place of each 3-second movie we presented three different still images taken from the beginning, middle and end of the corresponding movie clip. Each image was presented for one second with no ISI, to equate the total presentation time with the corresponding dynamic movie clip (Figure 1).

**Figure 1.** Examples of the static images taken from the 3-s movie clips depicting faces, bodies, scenes, objects, and scrambled objects. Still images taken from the beginning, middle and end of the corresponding movie clip.

In Experiment 3 visual field responses in face-selective regions were mapped using 2-sec video clips of dynamic faces making one of four different facial expressions: happy, fear, disgust and neutral air-puff. These faces were used in a previous fMRI study of face perception (van der Gaag et al., 2007). Happy expressions were recorded when actors laughed spontaneously at jokes, whereas the fearful and disgusted expressions were posed by the actors. The neutral, air-puff condition consisted of the actors blowing out their cheeks to produce movement but expressing no emotion. Both male and female actors were used. Videos were filmed against a grey background and the actors limited their head movements. Face videos were presented in the contralateral and ipsilateral visual hemifields at 5 by 5 degrees of visual angle and shown at a distance of 5 degrees from fixation to the edge of the stimulus (Pitcher et al., 2020).

### Procedure and Data Acquisition

#### Experiment 1 – Localizing the face-selective area in the lateral prefrontal cortex

Functional runs presented movie clips from five different stimulus categories (faces, bodies, scenes, objects, or scrambled objects). Data were acquired over 6 blocked-design functional runs lasting 234 seconds each. Each functional run contained three 18-second rest blocks, at the beginning, middle, and end of the run, during which a series of six uniform color fields were presented for three seconds each. Participants were instructed to watch the movies but were not asked to perform any overt task.

Each run contained two sets of five consecutive stimulus blocks (faces, bodies, scenes, objects, or scrambled objects) sandwiched between these rest blocks, to make two blocks per stimulus category per run. Each block lasted 18 seconds and contained stimuli from one of the five stimulus categories. The order of stimulus category blocks in each run was palindromic (e.g., fixation, faces, objects, scenes, bodies, scrambled objects, fixation, scrambled objects, bodies, scenes, objects, faces, fixation) and was randomized across runs.

Imaging data were collected using a 3T GE HDx Excite MRI scanner at the University of York. Functional images were acquired with an 8-channel phased array head coil (GE) and a gradient-echo EPI sequence (38 interleaved slices, repetition time (TR) = 3 sec, echo time (TE) = minimum full, flip angle =90%; voxel size 3mm isotropic; matrix size = 128 x 128) providing whole brain coverage. Slices were aligned with the anterior to posterior commissure line. Structural images were acquired using the same head coil and a high-resolution T-1 weighted 3D fast spoilt gradient (SPGR) sequence (176 interleaved slices, repetition time (TR) = 7.8 sec, echo time (TE) = minimum full, flip angle = 20 degrees; voxel size 1mm isotropic; matrix size = 256 × 256).

#### Experiment 2 – Measuring the response to moving and static stimuli in the right IFG

Functional data were acquired over 11 blocked-design functional runs lasting 234 seconds each. Each functional run contained three 18-second rest blocks, at the beginning, middle, and end of the run, during which a series of six uniform color fields were presented for three seconds. Participants were instructed to watch the movies and static images but were not asked to perform any overt task.

Functional runs presented either movie clips (the eight dynamic runs) or sets of static images taken from the same movies (the four static runs). For the dynamic runs, each 18-second block contained six 3-second movie clips from that category. For the static runs, each 18-second block contained 18 one-second still snapshots, composed of six triplets of snapshots taken at one second intervals from the same movie clip. Dynamic / static runs were run in the following order: 2 dynamic, 2 static, 2 dynamic, 2 static, 4 dynamic. The final 3 runs of the dynamic stimuli were used to define face-selective ROIs (see ‘Data Analysis’ section).

Imaging data were acquired using a 3T Siemens Magnetom Prisma MRI scanner (Siemens Healthcare, Erlangen, Germany) at the University of York. Functional images were acquired with a twenty-channel phased array head coil and a gradient-echo EPI sequence (38 interleaved slices, repetition time (TR) = 3 sec, echo time (TE) = minimum full, flip angle =90%; voxel size 3mm isotropic; matrix size = 128 x 128) providing whole brain coverage. Slices were aligned with the anterior to posterior commissure line. Structural images were acquired using the same head coil and a high-resolution T-1 weighted 3D fast spoilt gradient (SPGR) sequence (176 interleaved slices, repetition time (TR) = 7.8 sec, echo time (TE) = minimum full, flip angle = 20 degrees; voxel size 1mm isotropic; matrix size = 256 × 256).

#### Experiment 3 – Measuring the visual field response in the face area in the IFG

Participants fixated the center of the screen while 2-sec video clips of actors performing different facial expressions were shown in the four quadrants of the visual field. To ensure that participants maintained fixation, they were required to detect the presence of an upright or inverted letter (either a T or an L) at the center of the screen. Letters (0.6° in size) were presented at fixation for 250ms in random order and in different orientations at 4 Hz (Kastner et al., 1999). Participants were instructed to respond when the target letter (either T or L) was shown; this occurred approximately 25% of the time. The target letter (T or L) was alternated and balanced across participants. We informed the participants that the target detection task was the aim of the experiment, and we discarded any runs in which the participant scored less than seventy-five percent correct (Figure 2).

**Figure 2.** Static image taken from the hemifield visual field (VF) mapping stimulus used in Experiment 3. Actors displaying different emotions (happy, fear, disgust, neutral air-puff) were shown in the two hemifields. Participants maintained fixation by detecting the presence of either a T or an L (shown upright or inverted) at fixation (Pitcher et al., 2020).

Visual field mapping images were acquired over 6 blocked-design functional runs lasting 408 sec each. Each functional run contained sixteen 16-sec blocks during which eight videos of eight different actors performing the same facial expression (happy, fear, disgust and neutral air-puff) were presented in one of the two hemifields. Eight blocks were shown in each hemifield and the order in which they appeared was randomized. After the visual field mapping blocks were completed, participants completed 6 blocked-design functional runs lasting 234 sec each to functionally localize the face-selective ROIs.

### Brain Imaging Acquisition and Analysis

Imaging data were acquired using research dedicated GE 3-Tesla scanner at the National Institutes of Health (NIH). Functional images were acquired using with a 32-channel head coil and a gradient-echo EPI sequence (36 interleaved slices, repetition time (TR) = 2 sec, echo time (TE) = 30 ms; voxel size 3mm isotropic; 0.6 mm interslice gap) providing whole brain coverage. Slices were aligned with the anterior/posterior commissures. In addition, a high-resolution T-1 weighted MPRAGE anatomical scan (T1-weighted FLASH, 1 × 1 × 1 mm resolution) was acquired to anatomically localize functional activations.

### Imaging Analysis

Functional MRI data were analyzed using AFNI (http://afni.nimh.nih.gov/afni). Images were slice-time corrected and realigned to the third volume of the first functional run and to the corresponding anatomical scan. All data were motion corrected and any TRs in which a participant moved more than 0.3mm in relation to the previous TR were discarded from further analysis. The volume-registered data were spatially smoothed with a 4-mm full-width-half-maximum Gaussian kernel. Signal intensity was normalized to the mean signal value within each run and multiplied by 100 so that the data represented percent signal change from the mean signal value before analysis.

In Experiment 1 data from all six runs was entered into a general linear model (GLM) by convolving the standard hemodynamic response function with the regressors of interest (faces, bodies, scenes, objects and scrambled objects. Regressors of no interest (e.g., 6 head movement parameters obtained during volume registration and AFNI’s baseline estimates) were also included in the GLM. Data from all thirty-two participants were entered in a group whole brain analysis to identify the locus of the face-selective activations in the bilateral frontal cortex using a contrast of moving faces greater than moving objects (Figure 3).

**Figure 3.**
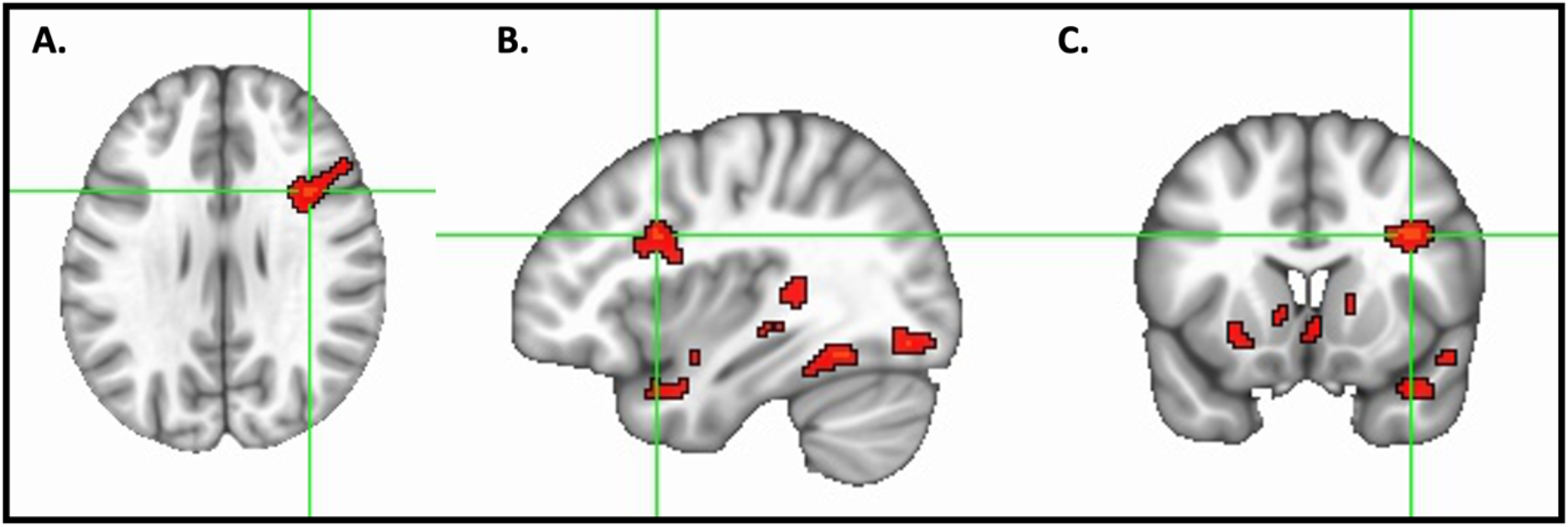
Results of a whole-brain group analysis (*N*=32) showing a contrast of moving faces greater than moving objects activations in red (t-statistical threshold is *p* =0.001, cluster correction of 50 voxels). The face-selective area in the right lateral prefrontal cortex is shown in the green crosshairs of a horizontal brain slice (Figure 3a), a sagittal brain slice (Figure 3b) and a coronal brain slice (Figure 3c). The peak voxel (MNI co-ordinates 37, 13, 28) was centred in the pars opercularis of the right inferior frontal gyrus (IFG) according to the probabilistic maps for combining functional imaging data with cytoarchitectonic maps (Eickhoff et al., 2005).

We next analyzed data for all participants individually to localize the regions of interest (ROIs). Face-selective ROIs were identified for each participant using a contrast of greater activation evoked by faces than that evoked by objects, calculating significance maps of the brain using an uncorrected statistical threshold of *p* = 0.001. In addition to the face-selective area in the prefrontal cortex we also identified the FFA, pSTS and OFA. Finally, we performed a split-half analysis to calculate the neural response to different stimulus categories (faces, bodies, scenes, objects and scrambled objects) in the face-selective ROIs. Even runs (2, 4 and 6) were used to identify the face-selective areas; odd runs (1, 3 and 5) were used to calculate the neural responses. Within each functionally defined ROI, we then calculated the magnitude of response (percent signal change from a fixation baseline) for each stimulus category. We selected all contiguous voxels for each ROI.

Data in Experiment 2 were analyzed using the same preprocessing procedures described in Experiment 1 except for the following differences. ROIs were calculated using data from 4 dynamic runs (runs 9 to 11). Face-selective ROIs were identified using a contrast of moving faces greater than moving objects using an uncorrected statistical threshold of *p* = 0.001. Within ROIs we then calculated the magnitude of response to the dynamic and static conditions of each of the five stimulus categories (faces, bodies, scenes, objects and scrambled objects), using the data collected from runs 1 to 8 in which pairs of dynamic and static runs were alternated. All the data used to calculate percent signal change (PSC) was independent of the data used to define the ROIs.

Data in Experiment 3 were analyzed using the same preprocessing procedures described in Experiment 1 except for the following differences. Face-selective ROIs were identified using data from 6 dynamic runs (7 to 12) using a contrast of moving faces greater than moving objects using an uncorrected statistical threshold of *p* = 0.001. Within ROIs we then calculated the magnitude of response to moving face videos presented in the contralateral and ipsilateral visual fields using the data collected from runs 1 to 6.

## Results

### Experiment 1: Localizing the face-selective area in the lateral prefrontal cortex

Data from all thirty-two participants were entered into a group whole brain ANOVA to identify the locus of the face-selective activations in the bilateral prefrontal cortex. The results of a contrast of moving faces greater than moving objects is shown in Figure 3. Using a t-statistical threshold of *p* = 0.001 and a cluster correction of 50 voxels we were able to localise a face-selective activation in the right lateral prefrontal cortex, but not in the left lateral prefrontal cortex. The face-selective activation in the right lateral prefrontal cortex was centred in the pars opercularis of the inferior frontal gyrus (MNI coordinates 37, 13, 28) according to the probabilistic maps for combining functional imaging data with cytoarchitectonic maps (Eickhoff et al., 2005). The activation was also within 1mm of the right inferior frontal junction where face-selective activation has previously been reported by other groups (Chan & Downing, 2011; Gobbini et al., 2004; Keightley et al., 2011).

To further characterise how reliable this activation was across all thirty-two participants we next looked at the individual level using data collected from all six experimental runs. Results revealed that a face-selective area in the frontal cortex was present in twenty-four participants in the right hemisphere (mean MNI co-ordinates 42, 14, 32), but only eighteen in the left hemisphere (mean MNI co-ordinates −38, 17, 33). By contrast we were able to localise the right FFA (mean MNI co-ordinates 41, −52, −17), left FFA (mean MNI co-ordinates −41, −52, −17), right pSTS (mean MNI co-ordinates 53, −37, 5) and left pSTS (mean MNI co-ordinates −57, −39, 6) in thirty-one of thirty-two participants. The right OFA was present in thirty participants (mean MNI co-ordinates 40, −79, −10) and the left OFA in twenty-five (mean MNI co-ordinates −39, −82, −10). These results demonstrate that the face-selective area in the IFG was not as reliably identified across participants as face-selective areas in the occipitotemporal cortex. This greater preference for face processing in the right hemisphere is consistent with prior evidence (Barton et al., 2002; Sliwinska & Pitcher, 2018; Young et al., 1985; Yovel et al., 2003).

Finally, to compare the response of the face-selective area in the middle frontal gyrus to other face-selective areas we performed a split-half analysis of our data (Figure 4). Because we were only able to localise the left IFG in eighteen of the participants we focused on the face-selective ROIs in the right hemisphere, but the responses in the left hemisphere ROIs showed the same overall pattern as those in the right hemisphere. We were able to identify the four ROIs of interest in twenty-two of the thirty-two participants (two participants who had face-selective activity in the IFG did not have a right OFA).

**Figure 4.**
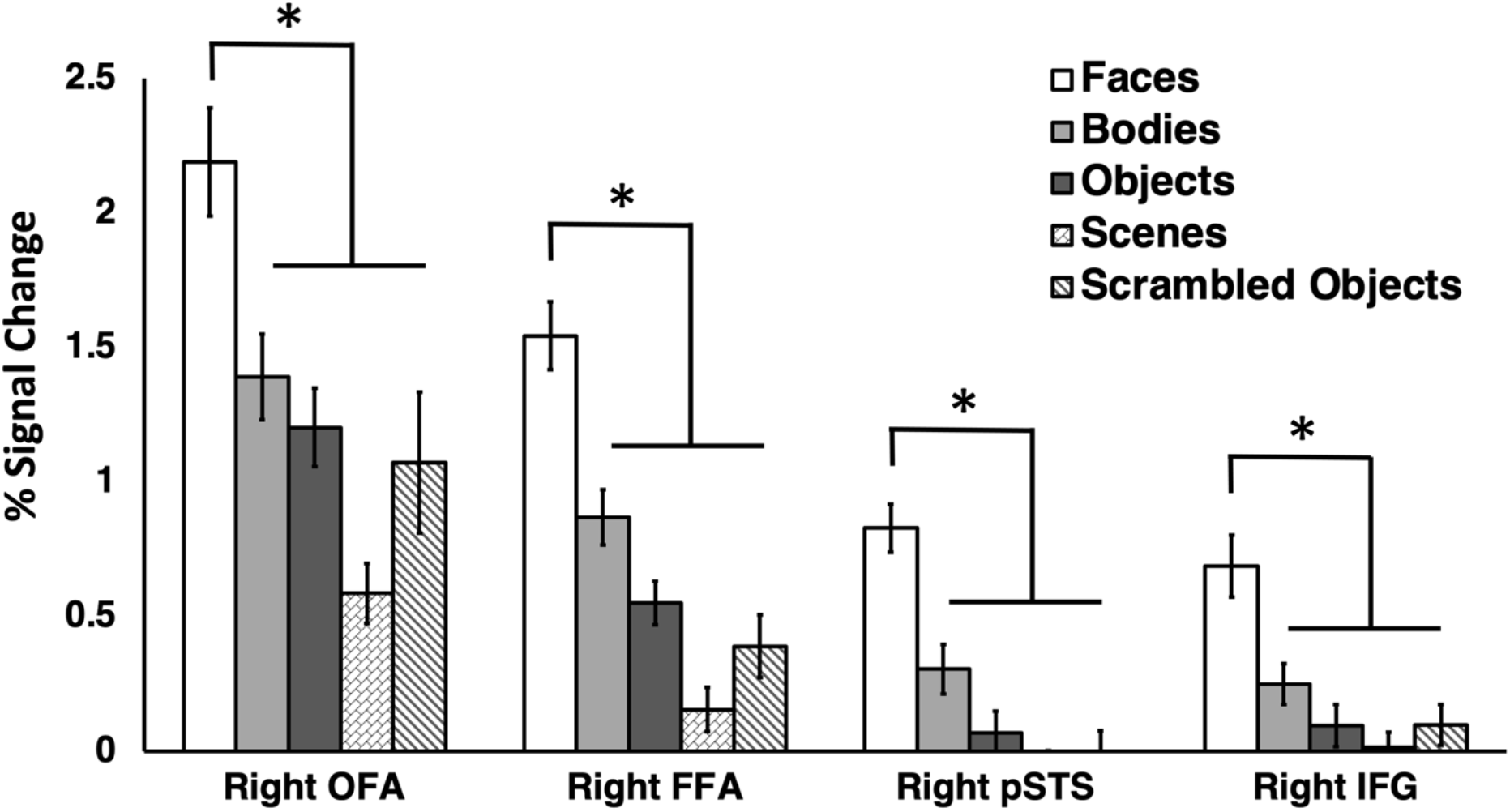
Percent signal change (PSC) data for the dynamic for five visual categories (faces, bodies, scenes, objects, and scrambled objects) in the rOFA, rFFA, rpSTS and rIFG. All four regions showed a significantly greater response to faces than all other categories. Data shown are independent of the data used to define the ROIs. Error bars show standard errors of the mean across participants. ∗ denotes a significant difference (*p* < 0.01) in post hoc tests.

Percent signal change data (Figure 4) were entered into a 2 (ROI: IFG, FFA, pSTS, OFA) by 5 (Category: Faces, Bodies, Scenes, Objects and Scrambled Objects) repeated measures analysis of variance (ANOVA). Results showed significant main effects of ROI (F (3,63) = 57, *p* < 0.001; partial η ^2^ = 0.731) and stimulus (F (4,84) = 72, *p* < 0.001; partial η ^2^ = 0.774). Stimulus and ROI also combined in a significant interaction (F (12,252) = 7.6, *p* < 0.001; partial η ^2^ = 0.265). Planned Bonferroni comparisons demonstrated that all four ROIs exhibited a significantly greater response to faces than to all other stimulus categories (*p* < 0.001).

Experiment 2 – Measuring the response to moving and static stimuli in the rIFG Face-selective ROIs were identified in both hemispheres using a contrast of moving faces greater than moving objects. As in Experiment 1 we were not able to localize face-selective ROIs in all twenty-four participants across both hemispheres. Results revealed that a face-selective area in the frontal cortex was present in seventeen participants in the right hemisphere (mean MNI co-ordinates 43, 8, 39), but only fourteen in the left hemisphere (mean MNI co-ordinates −45, 2, 39). By contrast we were able to localise the right FFA (mean MNI co-ordinates 41, −54, −17), left FFA (mean MNI co-ordinates −41, −54, −1), right pSTS (mean MNI co-ordinates 54, −43, 9) and left pSTS (mean MNI co-ordinates −55, −40, 9) in all participants. The right OFA was present in twenty-two participants (mean MNI co-ordinates 41, −84, −9) and the left OFA in eighteen participants (mean MNI co-ordinates −40, −83, −11). We again focused our analysis on the ROIs in the right hemisphere but the pattern in the left hemisphere ROIs was consistent.

To establish which face-selective ROIs showed a differential response to moving and static stimuli, we analysed the data in a 2 (motion: moving, static) by 5 (stimulus: bodies, faces, objects, scenes, scrambled objects) by 4 (ROI: FFA, OFA, pSTS, IFG) repeated-measures analysis of variance (ANOVA). We found significant main effects of motion (F (1,16) = 23, *p* < 0.001; partial η ^2^ = 0.587), stimulus (F (4,64) = 112, *p* < 0.001; partial η ^2^ = 0.875) and ROI (F (3,48) = 58, *p* < 0.001; partial η ^2^ = 0.784). Motion and Stimulus combined in a significant interaction (F (4,64) = 5.7, p < 0.001; partial η ^2^ = 0.265). Motion and ROI combined in a significant interaction (F (3,48) = 3.8, p = 0.015; partial η ^2^ = 0.195). Stimulus and ROI combined in a significant interaction (F (12,192) = 25, *p* < 0.001; partial η ^2^ = 0.607). Most importantly motion, stimulus and ROI combined in a significant three-way interaction (F (12,192) = 2.9, *p* = 0.001; partial η ^2^ = 0.154). To further understand what factors were driving the significant effects, we then performed separate two-way ANOVAs on each face-selective ROI.

#### Right IFG

A 2 (motion) × 5 (stimulus) repeated-measures ANOVA showed a main effect of motion (*F* (1, 16) = 6.8, *p* = 0.019; partial η ^2^ = 0.299) with a significantly greater response to moving more than static stimuli (*p* = 0.003). There was also a main effect of stimulus (*F* (4, 64) = 9, *p* < 0.001; partial η ^2^ = 0.361) with a greater response to faces than to all other stimulus categories (*p* < 0.05). There was also a significant interaction between motion and stimulus (*F* (4, 64) = 2.5, *p* = 0.048; partial η ^2^ = 0.137). Planned Bonferroni comparisons revealed that moving faces produced a larger response than static faces (*p* < 0.001), but no other comparisons reached significance (*p* > 0. 15).

#### Right pSTS

A 2 (motion) × 5 (stimulus) repeated-measures ANOVA showed a main effect of motion (*F* (1, 16) = 6.1, *p* = 0.026; partial η ^2^ = 0.290) with a significantly greater response to moving more than static stimuli (*p* = 0.026). There was also a main effect of stimulus (*F* (4, 64) = 47, *p* < 0.001; partial η ^2^ = 0.759) with a greater response to faces than to all other stimulus categories (*p* < 0.001). There was also a significant interaction between motion and stimulus (*F* (4, 64) = 13.5, *p* < 0.001; partial η ^2^ = 0.474). Planned Bonferroni comparisons revealed that moving faces produced a larger response than static faces (*p* < 0.001) and that moving bodies produced a larger response than static bodies (*p* = 0.05), but no other comparisons reached significance (*p* = 1).

#### Right FFA

A 2 (motion) × 5 (stimulus) repeated-measures ANOVA showed a main effect of motion (*F* (1, 16) = 8.1, *p* = 0.012; partial η ^2^ = 0.351) with a significantly greater response to moving more than static stimuli (*p* = 0.012). There was also a main effect of stimulus (*F* (4, 64) = 61, *p* < 0.001; partial η ^2^ = 0.801) with a greater response to faces than to all other stimulus categories (*p* < 0.01). There was no significant interaction between motion and stimulus (*F* (4, 64) = 1.5, *p* = 0.2; partial η ^2^ = 0.094).

#### Right OFA

A 2 (motion) x 5 (stimulus) repeated-measures ANOVA showed a main effect of motion (*F* (1, 16) = 45, *p* < 0.001; partial η ^2^ = 0.751) with a significantly greater response to moving more than static stimuli (*p* = < 0.001). There was also a main effect of stimulus (*F* (4, 64) = 53, *p* < 0.001; partial η ^2^ = 0.778) with a greater response to faces than to all other stimulus categories (*p* < 0.01). There was also a significant interaction between motion and stimulus (*F* (4, 64) = 3.6, *p* = 0.01; partial η ^2^ = 0.195). Planned Bonferroni comparisons revealed that moving faces produced a larger response than static faces (*p* < 0.001), moving objects produced a larger response than static objects (*p* < 0.001), moving bodies produced a larger response than static bodies (*p* < 0.001) and that moving scrambled objects produced a larger response than static scrambled objects (*p* = 0.01). There was no significant difference between moving and static scenes (*p* = 0.15).

### Experiment 3 – fMRI mapping of faces in the two hemifields in face-selective areas

Face-selective ROIs were identified in both hemispheres using a contrast of moving faces greater than moving objects. As in Experiment 1 we were not able to localize face-selective ROIs in all eighteen participants across both hemispheres. Results revealed that a face-selective area in the frontal cortex was present in sixteen participants in the right hemisphere (mean MNI co-ordinates 40, 10, 32), but only eleven in the left hemisphere (mean MNI co-ordinates −43, 15, 30). We again focused our analysis on the ROIs in the right hemisphere but the pattern in the left hemisphere ROIs was consistent.

To establish which face-selective ROIs showed a greater response to faces in the contralateral visual field we analysed the data in a 2 (visual field: ipsilateral, contralateral) by 4 (ROI: FFA, OFA, pSTS, IFG) repeated-measures analysis of variance (ANOVA). We found significant main effects of visual field (F (1,15) = 30, *p* < 0.001; partial η ^2^ = 0.669) with a significantly greater response to faces in the contralateral more than the ipsilateral visual field (*p* < 0.001). There was no main effect of ROI (F (3,45) = 1.8, *p* = 0.17; partial η ^2^ = 0.105). Importantly visual field and ROI combined in a significant two-way interaction (F (3,45) = 31, *p* < 0.001; partial η ^2^ = 0.671). Planned Bonferroni comparisons revealed a larger response to faces in the contralateral more than ipsilateral field in the FFA (*p* < 0.001) and OFA (*p* < 0.001) but not in the pSTS (*p* = 0.5) or IFG (*p* = 0.3).

## Discussion

The aim of the current study was to measure the response to visually presented images of faces in the human lateral prefrontal cortex and to compare these responses with those recorded in the face-selective areas in the occipitotemporal cortex (FFA, pSTS and OFA). In Experiment 1 we scanned thirty-two participants with fMRI while viewing short movie clips of faces, bodies, scenes, objects, and scrambled objects. Using a contrast of faces greater than objects we identified a face-selective area centred in the pars opercularis of the right inferior frontal gyrus (IFG), a finding consistent with prior fMRI studies (Chan & Downing, 2011; Gobbini et al., 2004; Keightley et al., 2011). A subsequent ROI analysis of individual participants revealed that this face-selective activation was present in only twenty-four of the thirty-two participants in the right hemisphere and in eighteen participants in the left hemisphere. By contrast, the bilateral FFA and pSTS areas were present in all but one of the participants, the right OFA was present in thirty participants and the left OFA in twenty-five. Even though the face area in the right IFG was less robustly identified across participants, it still exhibited the highly selective response to faces observed in the FFA, pSTS and OFA (Figure 4). In Experiment 2 we measured the response to moving and static stimuli across the four face-selective ROIs in the right hemisphere. Results demonstrated that the right IFG, right pSTS and right OFA all exhibited a greater response to moving faces than to static faces, but the right FFA responded equally to moving and static faces (Figure 5). Finally in Experiment 3 we measured responses to moving faces presented in the contralateral and ipsilateral visual fields. Results demonstrated the contralateral visual field bias observed in the right FFA and right OFA was absent in the right pSTS and right IFG (Figure 6). Taken together, the results of all three experiments suggest two principal conclusions. Firstly, that the face-selective area in the IFG is less robustly identified than face areas in the occipitotemporal cortex, this was observed in all three experiments. Secondly, that the similarity of the response patterns in the IFG and pSTS (greater response to moving faces more than static faces and no visual field bias) suggests that the two areas are functionally connected.

**Figure 5.**
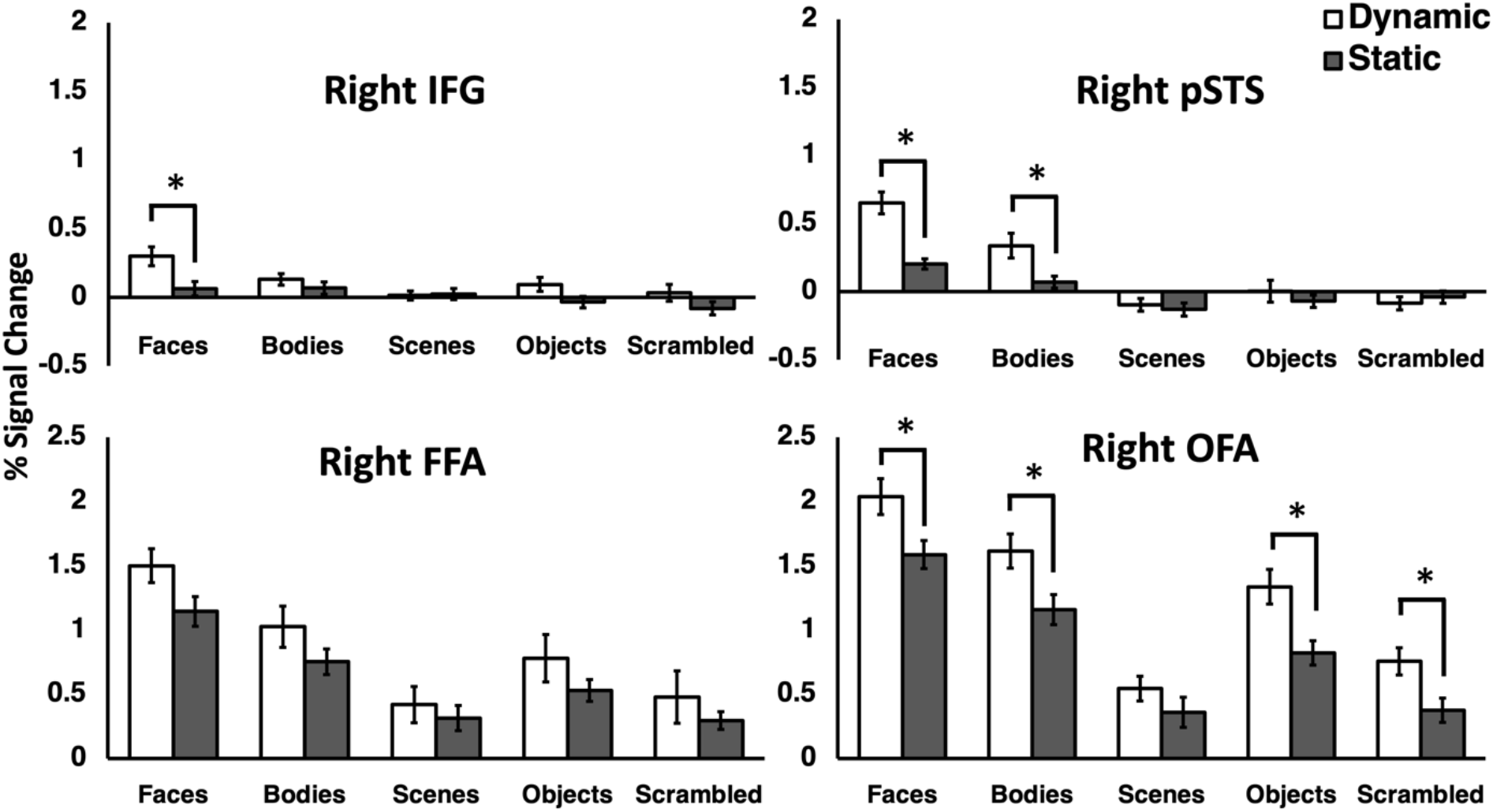
Percent signal change (PSC) data for the dynamic and static stimuli from all five categories (faces, bodies, scenes, objects and scrambled objects) in the IFG, rpSTS, rFFA and rOFA. All four regions showed a significantly greater response to faces than all other categories. The rIFG showed a greater response to moving faces than static faces. The rpSTS showed a greater response to moving faces than static faces and to moving bodies more than static bodies. The rOFA showed a greater response to moving more than static stimuli for four of the visual categories (face, bodies, objects, and scrambled objects). There was no significant difference moving and static stimuli in the rFFA. Error bars show standard errors of the mean across participants. ∗ denotes a significant difference (P < 0.0001) in post hoc tests.

**Figure 6.**
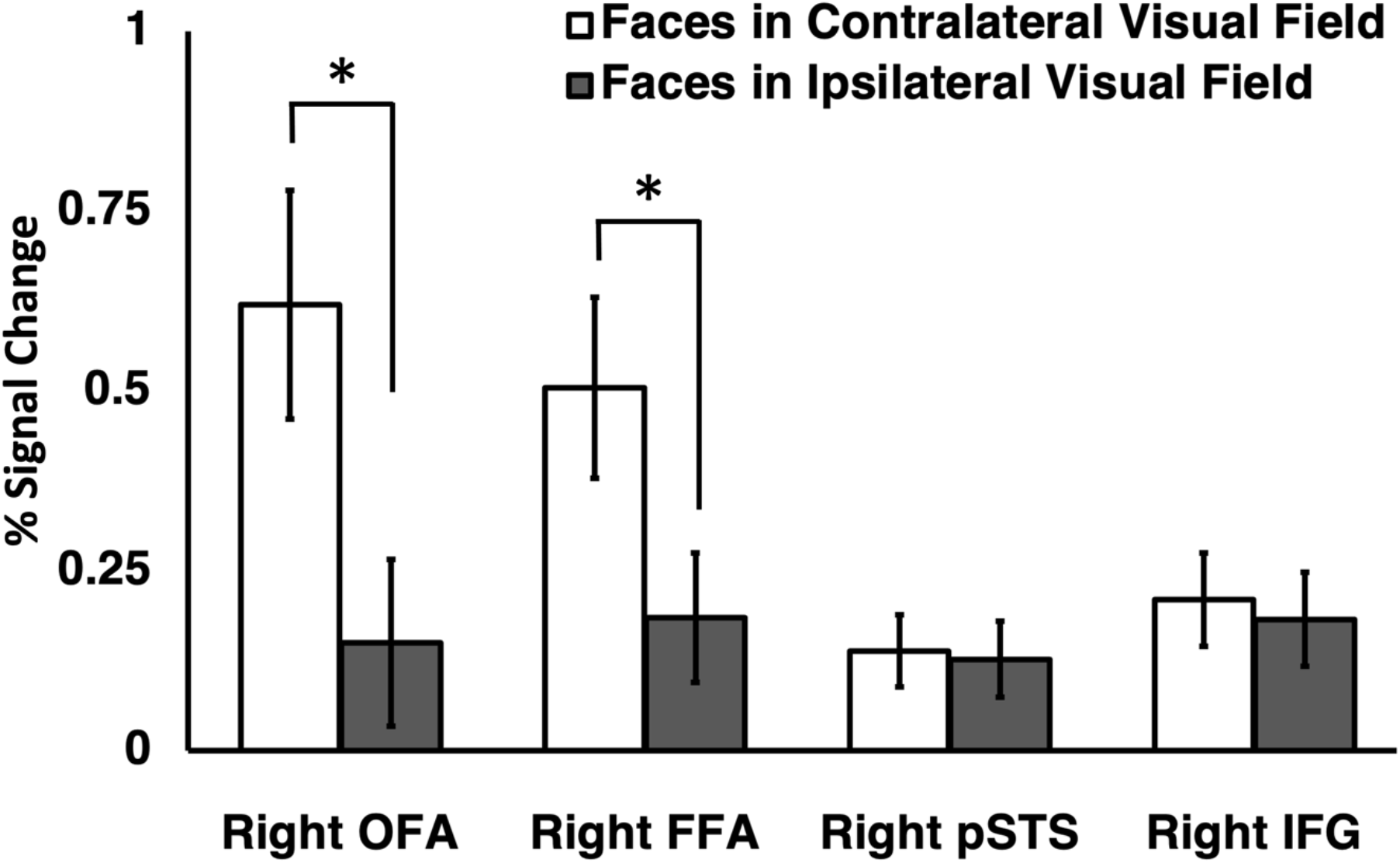
Percent signal change (PSC) for dynamic faces presented in the contralateral and ipsilateral hemifields. Results showed that the right FFA and right OFA exhibited a significantly greater response to faces in the contralateral VF than in the ipsilateral visual field. There were no visual field biases in the right pSTS or rright IFG. Error bars show standard errors of the mean across participants. ∗ denotes a significant difference (P < 0.0001) in post hoc tests.

Anatomical studies in non-human primates have identified a white matter pathway, that projects from the lateral superior temporal cortex into the inferior frontal cortex (Kravitz et al., 2011). In humans this pathway (the arcuate fasciculus) is more prominent than in non-human primates (Rilling et al., 2008) and is involved in a range of tasks including language (Dick & Tremblay, 2012) and face processing (Nakajima et al., 2018). A large-scale study of 680 participants further characterised this pathway using structural and functional connectivity data as a specialised sub-network with the wider face processing network (Wang et al., 2020). The authors further proposed that this sub-network is specialised for processing the dynamic and changeable aspects of faces that include recognising facial expressions and reading the intentions from a face. The results of the present study are consistent with this conclusion. In Experiment 2 we demonstrated that the right pSTS and right IFG both exhibited a greater response to moving faces than to static faces. We observed this pattern in our earlier fMRI study of moving and static faces, but we were only able to successfully localize the right IFG in seven of thirteen participants, so the result was not statistically warranted (Pitcher et al., 2011). The results of Experiment 2 resolve this issue.

Visual field mapping studies can be used to identify the functional connections between anatomically connected brain regions. For example, the functional and anatomical connections from primary visual cortex into the STS have been extensively mapped in macaques (Boussaoud et al., 1990; Desimone & Ungerleider, 1986; Ungerleider & Desimone, 1986). Our prior visual mapping studies in humans had demonstrated that the face-selective area in the pSTS lacked the contralateral visual field bias observed in the FFA and OFA (Pitcher et al., 2020; Sliwinska, Bearpark, et al., 2020) (see also (Finzi et al., 2021)). In the present study we re-analysed this earlier data and established that the face-selective area in the right IFG also exhibited no visual field bias (Figure 6). This result further suggests a functional connection between the face-selective areas in the pSTS and IFG. It is also likely the IRF is connected to our recently proposed third visual pathway for social perception (Pitcher et al., 2017; Pitcher & Ungerleider, 2021) but it should be noted the IFG is also connected to the dorsal visual pathway for action observation (Kilner, 2011).

The precise role of the IFG in memory processing, its lateralisation and whether it is object-specific, or domain general is unclear. Facial working memory, in which a representation of a face is maintained after it has been removed from view, activates prefrontal regions (Courtney et al., 1996, 1997). It has been proposed that right frontal activity may be associated with the maintenance of a simple, icon-like image of the face, whereas the left frontal activity represents a more elaborate face representation that is created after longer retention delays and is more easily maintained (Haxby et al., 1995). Regions in the frontal gyrus were found to be activated during visual imagery of faces but not during face perception (Ishai et al., 2002). During visual imagery, the frontal regions evoke top-down control for generating and maintaining visual images of faces. However, it is debated whether this process is category-selective and evokes different activation patterns in response to faces and objects (Mechelli et al., 2004) or not category-selective, as visual imagery of different objects evokes the same non-content related activity in the frontal cortex regardless of object category (Ishai et al., 2000). These studies provide evidence for the involvement of the prefrontal areas in cognitive control, working memory and perception. This suggests that these regions may represent a connection between top-down cognitive control processes and bottom-up perception and hence these areas may also be involved in familiarity judgement by comparing the internally stored information about a person to the perception of a face (Baldauf & Desimone, 2014; Heekeren et al., 2004). Neuropsychological evidence further support this hypothesis, as damage to the right prefrontal cortex causes false recognition, which is defined as the tendency to mistake unfamiliar faces for familiar ones without impairing other face-related processing (Rapcsak et al., 1996). False recognition in frontal patients is suggested to result from impaired strategic decision making and monitoring to determine whether a face is truly familiar, thus representing a control area between memory and perception.

The use of localizers in fMRI studies has been standard for over twenty years (Kanwisher et al., 1997; Saxe et al., 2006) but it is known that this approach does not always identify the necessary regions of interest (ROIs) in all participants (Duncan et al., 2009; Pitcher et al., 2011). In Experiment 1 we used six localizer runs to identify the face-selective ROIs, which was enough data to successfully localize the bilateral FFA, pSTS and right OFA in all participants. However, we were only able to identify the right IFG in twenty-four participants and the left IFG in eighteen participants. One possible explanation for this result is that we did not require subjects to perform any explicit task during the localizer runs (e.g., a one-back memory task). Such a task may not be necessary for identifying ROIs in high-level visual cortex but may be necessary for prefrontal cortex which is known to be engaged in cognitive functions like cognitive control (MacDonald et al., 2000), working memory (Curtis & D’Esposito, 2003), and Theory of Mind (Kalbe et al., 2010). It is therefore possible that future studies aiming to localise the face-selective area in the bilateral IFG should require participants to perform an explicit memory task in the localiser runs.

## Acknowledgements

Thanks to Nancy Kanwisher and Mbemba Jabbi for providing experimental stimuli.

## References

Baldauf, D., & Desimone, R. (2014) Neural mechanisms of object-based attention. Science, 344(6182), 424–427. https://doi.org/10.1126/science.1247003

Barton, J. J. S., Press, D. Z., Keenan, J. P., & O’Connor, M. (2002) Lesions of the fusiform face area impair perception of facial configuration in prosopagnosia [Article]. Neurology, 58(1), 71–78. https://doi.org/10.1212/WNL.58.1.71

Boussaoud, D., Ungerleider, L. G., & Desimone, R. (1990) Pathways for motion analysis: Cortical connections of the medial superior temporal and fundus of the superior temporal visual areas in the macaque [Article]. Journal of Comparative Neurology, 296(3), 462–495. https://doi.org/10.1002/cne.902960311

Calder, A. J., & Young, A. W. (2005) Understanding the recognition of facial identity and facial expression [Review]. Nature Reviews Neuroscience, 6(8), 641–651. https://doi.org/10.1038/nrn1724

Chan, A. W., & Downing, P. E. (2011) Faces and eyes in human lateral prefrontal cortex. Frontiers in human neuroscience, 5, 51.

Courtney, S. M., Ungerleider, L. G., Keil, K., & Haxby, J. V. (1996) Object and spatial visual working memory activate separate neural systems in human cortex. Cerebral cortex, 6(1), 39–49.

Courtney, S. M., Ungerleider, L. G., Keil, K., & Haxby, J. V. (1997) Transient and sustained activity in a distributed neural system for human working memory. Nature, 386(6625), 608–611.

Curtis, C. E., & D’Esposito, M. (2003) Persistent activity in the prefrontal cortex during working memory. Trends in cognitive sciences, 7(9), 415–423.

Davies-Thompson, J., & Andrews, T. J. (2012) Intra-and interhemispheric connectivity between face-selective regions in the human brain. Journal of Neurophysiology, 108(11), 3087–3095.

Desimone, R., & Ungerleider, L. G. (1986) Multiple visual areas in the caudal superior temporal sulcus of the macaque [Article]. Journal of Comparative Neurology, 248(2), 164–189. https://doi.org/10.1002/cne.902480203

Dick, A. S., & Tremblay, P. (2012) Beyond the arcuate fasciculus: consensus and controversy in the connectional anatomy of language. Brain, 135(Pt 12), 3529–3550. https://doi.org/10.1093/brain/aws222

Duncan, K. J., Pattamadilok, C., Knierim, I., & Devlin, J. T. (2009) Consistency and variability in functional localisers [Article]. NeuroImage, 46(4), 1018–1026. https://doi.org/10.1016/j.neuroimage.2009.03.014

Eickhoff, S. B., Stephan, K. E., Mohlberg, H., Grefkes, C., Fink, G. R., Amunts, K., & Zilles, K. (2005) A new SPM toolbox for combining probabilistic cytoarchitectonic maps and functional imaging data. NeuroImage, 25(4), 1325–1335. https://doi.org/10.1016/j.neuroimage.2004.12.034

Finzi, D., Gomez, J., Nordt, M., Rezai, A. A., Poltoratski, S., & Grill-Spector, K. (2021) Differential spatial computations in ventral and lateral face-selective regions are scaffolded by structural connections. Nat Commun, 12(1), 2278. https://doi.org/10.1038/s41467-021-22524-2

Fox, C. J., Iaria, G., & Barton, J. J. S. (2009) Defining the face processing network: Optimization of the functional localizer in fMRI [Article]. Human Brain Mapping, 30(5), 1637–1651. https://doi.org/10.1002/hbm.20630

Gauthier, I., Tarr, M. J., Moylan, J., Skudlarski, P., Gore, J. C., & Anderson, A. W. (2000) The fusiform ‘face area’ is part of a network that processes faces at the individual level [Article]. Journal of Cognitive Neuroscience, 12(3), 495–504. https://doi.org/10.1162/089892900562165

Gobbini, M. I., Leibenluft, E., Santiago, N., & Haxby, J. V. (2004) Social and emotional attachment in the neural representation of faces. NeuroImage, 22(4), 1628–1635. https://doi.org/10.1016/j.neuroimage.2004.03.049

Grill-Spector, K., Knouf, N., & Kanwisher, N. (2004) The fusiform face area subserves face perception, not generic within-category identification [Article]. Nature Neuroscience, 7(5), 555–562. https://doi.org/10.1038/nn1224

Haxby, J. V., Hoffman, E. A., & Gobbini, M. I. (2000) The distributed human neural system for face perception. Trends in cognitive sciences, 4(6), 223–233.

Haxby, J. V., Ungerleider, L. G., Horwitz, B., Maisog, J. M., Rapoport, S. I., & Grady, C. L. (1996) Face encoding and recognition in the human brain. Proceedings of the National Academy of Sciences, 93(2), 922–927.

Haxby, J. V., Ungerleider, L. G., Horwitz, B., Rapoport, S. I., & Grady, C. L. (1995) Hemispheric differences in neural systems for face working memory: A PET-rCBF study. Human Brain Mapping, 3(2), 68–82.

Heekeren, H. R., Marrett, S., Bandettini, P. A., & Ungerleider, L. G. (2004) A general mechanism for perceptual decision-making in the human brain. Nature, 431(7010), 859–862. https://doi.org/10.1038/nature02966

Heller, A. S., Johnstone, T., Shackman, A. J., Light, S. N., Peterson, M. J., Kolden, G. G., Kalin, N. H., & Davidson, R. J. (2009) Reduced capacity to sustain positive emotion in major depression reflects diminished maintenance of fronto-striatal brain activation. Proc Natl Acad Sci U S A, 106(52), 22445–22450. https://doi.org/10.1073/pnas.0910651106

Hemond, C. C., Kanwisher, N. G., & Op de Beeck, H. P. (2007) A preference for contralateral stimuli in human object- and face-selective cortex [Article]. PLoS ONE, 2(6), Article e574. https://doi.org/10.1371/journal.pone.0000574

Ishai, A., Haxby, J. V., & Ungerleider, L. G. (2002) Visual imagery of famous faces: effects of memory and attention revealed by fMRI. Neuroimage, 17(4), 1729–1741.

Ishai, A., Schmidt, C. F., & Boesiger, P. (2005) Face perception is mediated by a distributed cortical network. Brain research bulletin, 67(1-2), 87–93.

Ishai, A., Ungerleider, L. G., & Haxby, J. V. (2000) Distributed neural systems for the generation of visual images. Neuron, 28(3), 979–990.

Kalbe, E., Schlegel, M., Sack, A. T., Nowak, D. A., Dafotakis, M., Bangard, C., Brand, M., Shamay-Tsoory, S., Onur, O. A., & Kessler, J. (2010) Dissociating cognitive from affective theory of mind: a TMS study. cortex, 46(6), 769–780.

Kanwisher, N., McDermott, J., & Chun, M. M. (1997) The fusiform face area: A module in human extrastriate cortex specialized for face perception. Journal of Neuroscience, 17(11), 4302–4311. https://doi.org/10.1523/jneurosci.17-11-04302.1997

Kay, K. N., Weiner, K. S., & Grill-Spector, K. (2015) Attention reduces spatial uncertainty in human ventral temporal cortex ventral temporal cortex. Curr Biol, 25, 1–6.

Keightley, M. L., Chiew, K. S., Anderson, J. A., & Grady, C. L. (2011) Neural correlates of recognition memory for emotional faces and scenes. Soc Cogn Affect Neurosci, 6(1), 24–37. https://doi.org/10.1093/scan/nsq003

Kilner, J. M. (2011) More than one pathway to action understanding [Review]. Trends in Cognitive Sciences, 15(8), 352–357. https://doi.org/10.1016/j.tics.2011.06.005

Kravitz, D. J., Saleem, K. S., Baker, C. I., & Mishkin, M. (2011) A new neural framework for visuospatial processing [Review]. Nature Reviews Neuroscience, 12(4), 217–230. https://doi.org/10.1038/nrn3008

LaBar, K. S., Crupain, M. J., Voyvodic, J. T., & McCarthy, G. (2003) Dynamic perception of facial affect and identity in the human brain [Article]. Cerebral Cortex, 13(10), 1023–1033. https://doi.org/10.1093/cercor/13.10.1023

Leveroni, C. L., Seidenberg, M., Mayer, A. R., Mead, L. A., Binder, J. R., & Rao, S. M. (2000) Neural systems underlying the recognition of familiar and newly learned faces. Journal of Neuroscience, 20(2), 878–886.

Li, J., Liu, J., Liang, J., Zhang, H., Zhao, J., Huber, D. E., Rieth, C. A., Lee, K., Tian, J., & Shi, G. (2009) A distributed neural system for top-down face processing. Neuroscience letters, 451(1), 6–10.

MacDonald, A. W., Cohen, J. D., Stenger, V. A., & Carter, C. S. (2000) Dissociating the role of the dorsolateral prefrontal and anterior cingulate cortex in cognitive control. Science, 288(5472), 1835–1838.

Mechelli, A., Price, C. J., Friston, K. J., & Ishai, A. (2004) Where bottom-up meets top-down: neuronal interactions during perception and imagery. Cerebral cortex, 14(11), 1256–1265.

Nakajima, R., Yordanova, Y. N., Duffau, H., & Herbet, G. (2018) Neuropsychological evidence for the crucial role of the right arcuate fasciculus in the face-based mentalizing network: A disconnection analysis. Neuropsychologia, 115, 179–187. https://doi.org/10.1016/j.neuropsychologia.2018.01.024

Parvizi, J., Jacques, C., Foster, B. L., Witthoft, N., Rangarajan, V., Weiner, K. S., & Grill-Spector, K. (2012) Electrical stimulation of human fusiform face-selective regions distorts face perception. J Neurosci, 32(43), 14915–14920. https://doi.org/10.1523/JNEUROSCI.2609-12.2012

Phillips, M. L., Young, A. W., Scott, S. K., Calder, A. J., Andrew, C., Giampietro, V., Williams, S. C. R., Bullmore, E. T., Brammer, M., & Gray, J. A. (1998) Neural responses to facial and vocal expressions of fear and disgust [Article]. Proceedings of the Royal Society B: Biological Sciences, 265(1408), 1809–1817,1851. https://doi.org/10.1098/rspb.1998.0506

Pilz, K. S., Bülthoff, H. H., & Vuong, Q. C. (2009) Learning influences the encoding of static and dynamic faces and their recognition across different spatial frequencies [Article]. Visual Cognition, 17(5), 716–735. https://doi.org/10.1080/13506280802340588

Pitcher, D., Dilks, D. D., Saxe, R. R., Triantafyllou, C., & Kanwisher, N. (2011) Differential selectivity for dynamic versus static information in face-selective cortical regions [Article]. NeuroImage, 56(4), 2356–2363. https://doi.org/10.1016/j.neuroimage.2011.03.067

Pitcher, D., Duchaine, B., & Walsh, V. (2014) Combined TMS and fMRI reveal dissociable cortical pathways for dynamic and static face perception [Article]. Current Biology, 24(17), 2066–2070. https://doi.org/10.1016/j.cub.2014.07.060

Pitcher, D., Ianni, G., & Ungerleider, L. G. (2019) A functional dissociation of face-, body-and scene-selective brain areas based on their response to moving and static stimuli [Article]. Scientific Reports, 9(1), Article 8242. https://doi.org/10.1038/s41598-019-44663-9

Pitcher, D., Japee, S., Rauth, L., & Ungerleider, L. G. (2017) The superior temporal sulcus is causally connected to the amygdala: A combined TBS-fMRI study [Article]. Journal of Neuroscience, 37(5), 1156–1161. https://doi.org/10.1523/JNEUROSCI.0114-16.2016

Pitcher, D., Pilkington, A., Rauth, L., Baker, C., Kravitz, D. J., & Ungerleider, L. G. (2020) The Human Posterior Superior Temporal Sulcus Samples Visual Space Differently from Other Face-Selective Regions [Article]. Cerebral Cortex, 30(2), 778–785. https://doi.org/10.1093/cercor/bhz125

Pitcher, D., & Ungerleider, L. G. (2021) Evidence for a Third Visual Pathway Specialized for Social Perception [Review]. Trends in Cognitive Sciences, 25(2), 100–110. https://doi.org/10.1016/j.tics.2020.11.006

Pitcher, D., Walsh, V., Yovel, G., & Duchaine, B. (2007) TMS evidence for the involvement of the right occipital face area in early face processing. Curr Biol, 17(18), 1568–1573. https://doi.org/10.1016/j.cub.2007.07.063

Puce, A., Allison, T., Bentin, S., Gore, J. C., & McCarthy, G. (1998) Temporal cortex activation in humans viewing eye and mouth movements [Article]. Journal of Neuroscience, 18(6), 2188–2199. https://doi.org/10.1523/jneurosci.18-06-02188.1998

Rapcsak, S. Z., Polster, M. R., Glisky, M. L., & Comers, J. F. (1996) False recognition of unfamiliar faces following right hemisphere damage: neuropsychological and anatomical observations. Cortex, 32(4), 593–611.

Renzi, C., Schiavi, S., Carbon, C.-C., Vecchi, T., Silvanto, J., & Cattaneo, Z. (2013) Processing of featural and configural aspects of faces is lateralized in dorsolateral prefrontal cortex: a TMS study. Neuroimage, 74, 45–51.

Rezlescu, C., Pitcher, D., & Duchaine, B. (2012) Acquired prosopagnosia with spared within-class object recognition but impaired recognition of degraded basic-level objects [Article]. Cognitive Neuropsychology, 29(4), 325–347. https://doi.org/10.1080/02643294.2012.749223

Rilling, J. K., Glasser, M. F., Preuss, T. M., Ma, X., Zhao, T., Hu, X., & Behrens, T. E. (2008) The evolution of the arcuate fasciculus revealed with comparative DTI. Nat Neurosci, 11(4), 426–428. https://doi.org/10.1038/nn2072

Rossion, B., Caldara, R., Seghier, M., Schuller, A. M., Lazeyras, F., & Mayer, E. (2003) A network of occipito-temporal face-sensitive areas besides the right middle fusiform gyrus is necessary for normal face processing [Article]. Brain, 126(11), 2381–2395. https://doi.org/10.1093/brain/awg241

Rotshtein, P., Henson, R. N. A., Treves, A., Driver, J., & Dolan, R. J. (2005) Morphing Marilyn into Maggie dissociates physical and identity face representations in the brain [Article]. Nature Neuroscience, 8(1), 107–113. https://doi.org/10.1038/nn1370

Saxe, R., Brett, M., & Kanwisher, N. (2006) Divide and conquer: A defense of functional localizers [Note]. NeuroImage, 30(4), 1088–1096. https://doi.org/10.1016/j.neuroimage.2005.12.062

Scalaidhe, S. P., Rodman, H. R., Albright, T. D., & Gross, C. G. (1997) The effects of combined superior temporal polysensory area and frontal eye field lesions on eye movements in the macaque monkey. Behav Brain Res, 84(1-2), 31–46. https://doi.org/10.1016/s0166-4328(96)00131-3

Schultz, J., Brockhaus, M., Bülthoff, H. H., & Pilz, K. S. (2013) What the human brain likes about facial motion [Article]. Cerebral Cortex, 23(5), 1167–1178. https://doi.org/10.1093/cercor/bhs106

Shepherd, S. V., & Freiwald, W. A. (2018) Functional Networks for Social Communication in the Macaque Monkey. Neuron, 99(2), 413–420 e413. https://doi.org/10.1016/j.neuron.2018.06.027

Sliwinska, M. W., Bearpark, C., Corkhill, J., McPhillips, A., & Pitcher, D. (2020) Dissociable pathways for moving and static face perception begin in early visual cortex: Evidence from an acquired prosopagnosic [Article]. Cortex, 130, 327–339. https://doi.org/10.1016/j.cortex.2020.03.033

Sliwinska, M. W., Elson, R., & Pitcher, D. (2020) Dual-site TMS demonstrates causal functional connectivity between the left and right posterior temporal sulci during facial expression recognition [Article]. Brain Stimulation, 13(4), 1008–1013. https://doi.org/10.1016/j.brs.2020.04.011

Sliwinska, M. W., & Pitcher, D. (2018) TMS demonstrates that both right and left superior temporal sulci are important for facial expression recognition [Article]. NeuroImage, 183, 394–400. https://doi.org/10.1016/j.neuroimage.2018.08.025

Tsao, D. Y., Moeller, S., & Freiwald, W. A. (2008) Comparing face patch systems in macaques and humans [Article]. Proceedings of the National Academy of Sciences of the United States of America, 105(49), 19514–19519. https://doi.org/10.1073/pnas.0809662105

Ungerleider, L. G., & Desimone, R. (1986) Cortical connections of visual area MT in the macaque [Article]. Journal of Comparative Neurology, 248(2), 190–222. https://doi.org/10.1002/cne.902480204

Wang, Y., Metoki, A., Smith, D. V., Medaglia, J. D., Zang, Y., Benear, S., Popal, H., Lin, Y., & Olson, I. R. (2020) Multimodal mapping of the face connectome. Nat Hum Behav, 4(4), 397–411. https://doi.org/10.1038/s41562-019-0811-3

Winston, J. S., Henson, R. N. A., Fine-Goulden, M. R., & Dolan, R. J. (2004) fMRI-adaptation reveals dissociable neural representations of identity and expression in face perception [Article]. Journal of Neurophysiology, 92(3), 1830–1839. https://doi.org/10.1152/jn.00155.2004

Young, A. W., Hay, D. C., McWeeny, K. H., Ellis, A. W., & Barry, C. (1985) Familiarity decisions for faces presented to the left and right cerebral hemispheres [Article]. Brain and Cognition, 4(4), 439–450. https://doi.org/10.1016/0278-2626(85)90032-6

Yovel, G., Levy, J., Grabowecky, M., & Paller, K. A. (2003) Neural correlates of the left-visual-field superiority in face perception appear at multiple stages of face processing [Article]. Journal of Cognitive Neuroscience, 15(3), 462–474. https://doi.org/10.1162/089892903321593162

